# Melanogenic Activity Facilitates Dendritic Cell Maturation via FMOD

**DOI:** 10.1101/2022.05.14.491976

**Authors:** Marianna Halasi, Irit Adini

**Author notes:** Correspondence to: Irit Adini, Department of Surgery, Harvard Medical School, Center for Engineering in Medicine & Surgery, Massachusetts General Hospital, 51 Bloosm Street, Boston, MA 02114 USA.

## Abstract

According to epidemiological research, autoimmune diseases are more prevalent among African Americans, therefore we hypothesized that pigment production in the microenvironment contribute to local immune regulation. Here, in an *in vitro* setting we examined the role for pigment production by murine epidermal melanocytes in immune and inflammatory responses via DC activation. Our results revealed that dark pigmented melanocytes increase the production of IL-3 and the pro-inflammatory cytokines IL-6 and TNF-a, and consequently they induce pDC maturation. Further, we found that low pigment associated FMOD interferes with cytokine secretion and subsequent pDC maturation. To the best of our knowledge, this is the first study to assess the effect of baseline pigmentation on epidermal melanocyte cytokine profile, and its impact on DCs.

## INTRODUCTION

Substantial number of epidemiological studies have demonstrated strong correlation between skin color and inflammatory diseases including but not limited to Lupus, Atopic Dermatitis, Neuromyelitis Optica, and autoimmune uveitis.^1–5^ These reports show undisputable disparities among racial and ethnic groups with respect to the severity, prevalence, and incidence rate of inflammation related disorders.

While black/brown and white people have a comparable number of melanocytes, their level of pigment production, as well as melanocyte structure and function are different.^6^ Melanocytes, the neural crest-derived melanin-producing cells^7^ are located in the basal layer of the epidermis within the extracellular matrix (ECM), where they comprise 3–5% of the total epidermal cell population; in addition, they can be found in hair bulbs, the retina, the inner ear, heart valves and septa, and meninges of the brain.^8,9^ Melanin (pigment) production takes place in lipid membrane covered organelles called melanosomes^10^ and is regulated by multiple factors such as hormones, auto-, para-, and intracrine signals via receptor-dependent and independent pathways.^11^ Melanogenesis, the biosynthesis of melanin (eumelanin-black/brown and pheomelanin-red/yellow) is mediated mainly by melanocyte-stimulating hormone (a-MSH) and adrenocorticotropic hormone (ACTH), after binding to their receptor, melanocortin 1 receptor (MC1R).^12^ Upon activation of the MC1R,which is highly polymorphic and affects the appearance of the skin tone, the level of intracellular cAMP increases, leading to elevated expression of melanogenic enzymes such as tyrosinase (TYR), mutations in TYR causes albinism, tyrosinase related protein 1 and 2 (TYRP1 and TYRP2).^13,14^

Besides protecting the skin from UV damage as their primary function, melanocytes are also scavengers of free radicals and have the capacity to reinforce the immune depositing cells that contribute to the innate immune response by activating skin-resident T cells like other antigen presenting cells (APCs).^15–17^ Moreover, numerous studies have shown that epidermal melanocytes are capable of phagocytosis, antigen processing and presentation, secretion of proinflammatory cytokines or immunosuppressive molecules in response to inflammatory stimuli. In addition, epidermal melanocytes act as non-professional APCs participating in local immune responses thus playing an important role in innate immunity.^18–27^ However, little is known about the role of pigmented melanocytes in stimulating inflammatory immune responses.

Dendritic cells (DCs), the most potent professional APCs play pivotal roles in the immunological homeostasis.^28^ In steady state conditions, when pathogens or inflammation are absent, DCs shape self-tolerance by presenting self-antigens to developing and mature T-cells.

Upon pathogen invasion, DCs are activated and initiate innate and adaptive immune responses by migrating to lymph nodes where they present antigens to effector T cells or interact with B cells.^29^ DCs differentiate in the bone marrow (BM) from hematopoietic stem cells (HSCs) via intermediate sequential progenitors.^30^ There are two main types of FMS-like tyrosine kinase 3 ligand (FLT3L)-dependent DCs originated from committed DC precursors (CDPs): conventional DCs (cDCs) and plasmacytoid DCs (pDCs).^31–33^ The cDC subtype found in peripheral and lymphoid tissues can be further categorized into cDC1 (CD8a^+^ -in lymphoid tissues; CD103^+^ - in non-lymphoid tissues) responsible for cross-presentation to CD8 T cells, and cDC2 subpopulations (CD11b+) inducing Th2 and Th17 immune responses as a result of priming CD4 T cells.^33^ The pDC subtype mainly resides in the blood and lymphoid tissues and is well-recognized for producing large amounts of type I interferons mainly upon viral but also during bacterial infection.^34–37^

The extracellular matrix (ECM), a complex network of proteins and proteoglycans produced by various cells, maintains the balance of tissue homeostasis. Fibromodulin (FMOD), a member of the class II small leucine-rich proteoglycan family plays a role in ECM organization.^38,39^ FMOD binds collagen thus facilitates the formation of collagen fibrils.^40^ Furthermore, it is involved in muscle development,^41^ cell reprogramming,^42^ angiogenesis^43–45^ and wound healing.^46^ Our prior work demonstrated that lowly pigmented melanocytes secrete high levels of FMOD and that FMOD increases TGF-β1 secretion.^44^ TGF-β1 has been described as an inhibitor of DC maturation.^47^

Here, we present evidence that melanocyte pigmentation and melanocyte-secreted factor, FMOD play important roles in the maturation of bone marrow derived (BM) DCs.

## METHODS

### Melanocytes, melanocyte conditioned media (CM) and treatment

Melan-C (non-pigmented C from BALB/c mice), Melan Cc (pigmented Cc isolated from BALB/c mice with a corrective point mutation of Tyrosinase by RNA-DNA; G-T at 291 oligonucleotide),^48^ Melan-e1 (lowly pigmented e1 from Black C57BL/6J mice with mutation on MC1R), and Melan-a (pigmented a from Black C57BL/6J mice),^49,50^ mouse melanocytes were grown in RPMI-1640 medium (Sigma) supplemented with 10% not-heat-inactivated fetal bovine serum (FBS) (Peak Serum), 1% penicillin-streptomycin(P/S) (GIBCO), 0.5% 2-mercaptoethanol (2-ME) (GIBCO) and 1% HEPES (GIBCO). FMOD-KD lowly pigmented Melan C and Melan e1 cells were generated with the Alt-R CRISPR -Cas9 system from IDT as per the recommendation of the manufacturer. In addition, Melan-C, Melan-Cc and Melan-a cells were grown with 200 nM tetradecanoyl phorbol acetate (TPA) (Sigma), and Melan-e1 cells with 200 nM TPA and 40 pM cholera toxin (Sigma). The cells were maintained at 37°C in 10% CO_2_. For the generation of conditioned medium (CM), equal numbers of darkly pigmented and lowly pigmented melanocytes, also control and FMOD-KD lowly pigmented melanocytes were plated and a day later the plates were rinsed with PBS and the growth media was replaced with media without the addition of TPA or 2-ME. For Forskolin treatment 20mM of Forskolin (Sigma) was added to the media and as for the control equal amount of dimethyl sulfoxide (DMSO) was added. Twenty-four hours later the CM was collected, and the melanocytes were counted. The amount of CM was calculated relative to 1-1.8 x 10^6^ cells/3 ml.

### Bone marrow-derived dendritic cell (BMDC) generation and treatment

Bone marrow of 8-week-old C57BL/6J male mice purchased from The Jackson Laboratory (Bar Harbor, ME, USA) was flushed from femurs and tibias with phosphate-buffered saline (PBS). The BM cells per animal were spined down and then resuspended in 1 mL Red Blood Cell Lysis (RBCL) buffer (Biolegend) at room temperature for 1 min to lyse the red blood cells. Then, 9 mL of growth medium (same as for melanocytes) was added to the cells, and they were centrifuged again. The BM cells per animal were resuspended in growth medium and cultured at 1-2 x 10^6^ cells/mL on sterile 6-well plates for 9 days in the presence of 200ng/ml recombinant murine Flt3-Ligand (Peprotech). The BM cells were maintained at 37°C in 5% CO_2_. On day 9, BMDCs in the nonadherent fraction were pulled, and the wells were rinsed with PBS, then the BMDCs were spined, counted, and equally distributed on fresh 6-well plates for incubation with the different CMs. Twenty-four hours later, both adherent and nonadherent BMDCs were collected and processed for flow cytometric analysis.

### Flow cytometry

The collected BMDCs were centrifuged once and resuspended in cell staining buffer (PBS supplemented with 2% FBS) containing anti-Fc receptor mAb (Biolegend). Cells were blocked for 10 min at 4°C, and then cell surface staining was performed for 30 min at 4°C with fluorochrome-conjugated monoclonal antibodies to mouse antigens purchased from Biolegend: XCR1, CD11b, B220, PDCA-1, CD40, CD80, CD86, MHC II. Following staining cells were washed and resuspended in staining buffer containing DAPI (Vector Laboratory) to differentiate between live and dead cells. Samples were analyzed by the FACSAria cell sorter (BD Biosciences), and the acquired data were analyzed using FlowJo software (version 10.6.1).

### Total RNA extraction and quantitative real-time PCR (qRT-PCR)

Melanocyte cells were processed by the IBI Isolate reagent (IBI Scientific) for total RNA extraction. The High-Capacity cDNA Reverse Transcription Kit (Applied Biosystems) was used to synthesize complementary DNA (cDNA). Quantitative real-time PCR reactions were run on the LightCycler 480 instrument (Roche) using the PrimeTime Gene Expression Master Mix (IDT DNA) and the following mouse primers: IL-3-S, 5’-CCT GCC TAC ATC TGC GAA T-3’, IL-3-AS, 5’-GAT CGT TAA GGT GGA CCA TGT-3; IL-6-S, 5’-AGC CAG AGT CCT TCA GAG A-3’, IL-6-AS, 5’-TCC TTA GCC ACT CCT TCT GT-3; TGF-b1-S, 5’-GCG GAC TAC TAT GCT AAA GAG G-3’, TGF-b1-AS, 5’-CCG AAT GTC TGA CGT ATT GAA GA-3; TNF-a-S, 5’-AGA CCC TCA CAC TCA GAT CA-3’, TNF-a-AS, 5’-TCT TTG AGA TCC ATG CCG TTG-3; PPIA-S, 5’-CAT CCT AAA GCA TAC GGG TCC-3’, PPIA-AS, 5’-TCT TTC ACT TTG CCA AAC ACC-3; GAPDH-S, 5’-AAT GGT GAA GGT CGG TGT G-3’, GAPDH-AS, 5’-GTG GAG TCA TAC TGG AAC ATG TAG-3’.

### Statistical analysis

Statistical analysis was performed with Graphpad as described in the figure legends. *P* values of <0.05 were considered to be statistically significant.

## RESULTS

### Dark pigmented melanocytes stimulate the maturation of the pDC subset of BMDCs

To explore a potential relationship between pigment production and maturation of dendritic cells, we investigated whether BMDCs could be activated by melanocytes. We utilized Flt3L to differentiate bone marrow cells into dendritic cells, as Flt3L supplemented cultures generate all three DC subsets (cDC1, cDC2 and pDC) with an immature phenotype resembling *in vivo* steady state conditions.^51,52^ Flt3L-BMDCs established from C57-black mice were incubated with growth media, conditioned medium (CM), of darkly pigmented (black: melan-a) and lowly pigmented (white: melan-e1) melanocytes for 24 hrs. As immature DCs (imDCs) undergo maturation they present increased levels of cellular reorganization and co-stimulatory markers, such as CD40, CD80, CD86 and major histocompatibility complex II (MHC II).^53^ Interestingly, regardless of melanocyte pigmentation the number of cDCs increased after exposure of BMDCs to CM of black and white melanocytes when compared to media control, but the number of pDCs were not affected (Suppl. Fig. 1A, top panel; Suppl. Fig. 1B). Although, CM of black melanocytes significantly stimulated the maturation of the pDC subset as assessed by the increased expression of cell surface maturation markers via flow cytometry when compared to CM of white melanocytes (Fig. 1A) and media control (Suppl. Fig. 1C), it did not affect notably the distribution of the DC subsets (Suppl. Fig. 1B). We did not observe difference in the degree of cDC maturation following incubation with CM of differentially pigmented melanocytes (data not shown).

**Figure 1.**
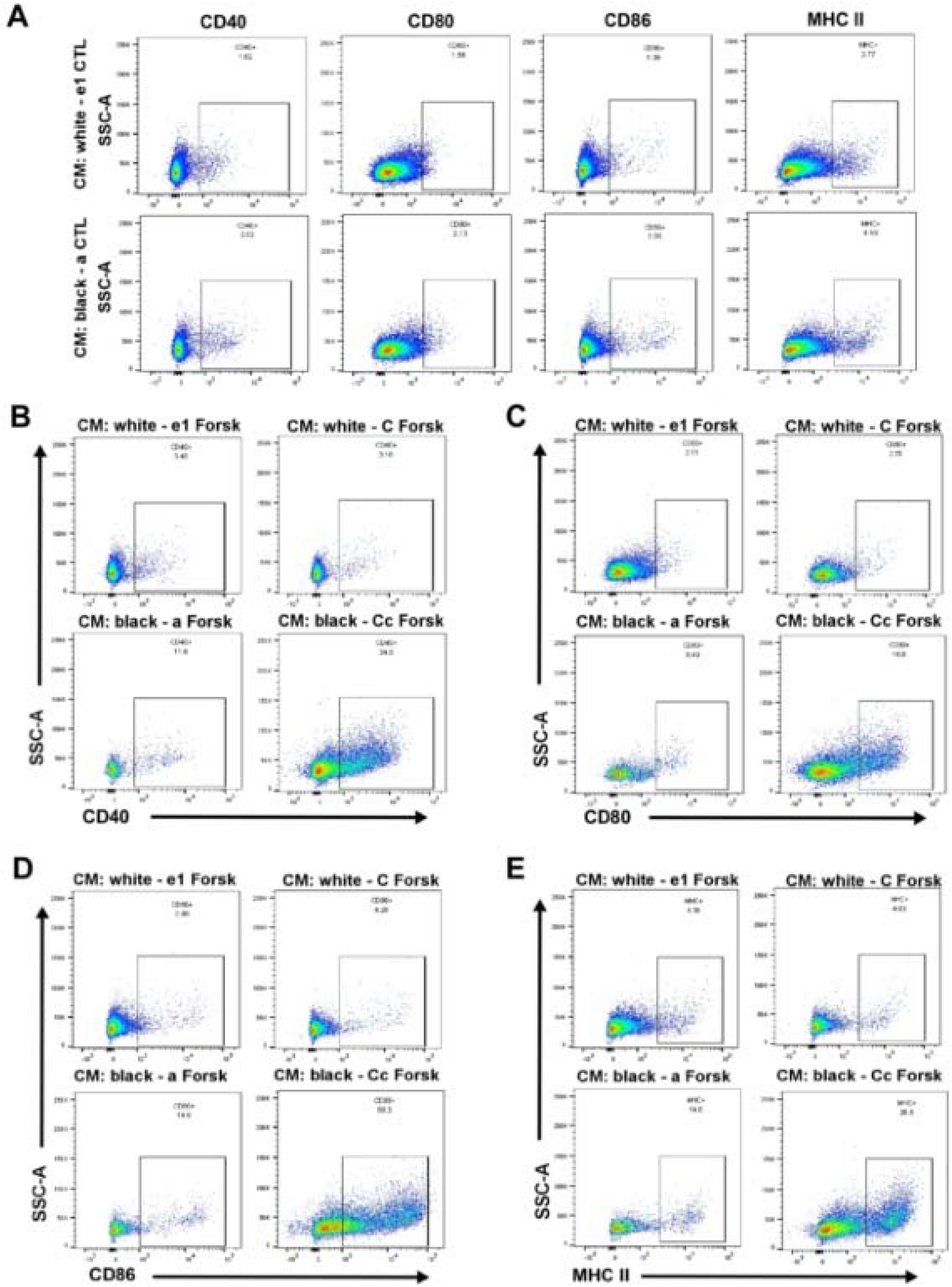
Dark pigmentation correlates with increased pDC maturation *ex vivo.* **(A)** Mouse Flt3L differentiated BM-DCs of 3 animals were incubated with CM derived from white (melan e1) and black (melan a) melanocytes. Twenty-four hours later the BM-DCs were analyzed for the maturity of the pDC subpopulation by flow cytometry with CD40, CD80, CD86 and MHC II surface markers. Representative data are shown as dot plots, and numbers represent the percentage of CD11b^-^XCR-1^-^PDCA-1^+^B220^+^ CD40^+^/CD80^+^/CD86^+^/MHC II^+^ pDCs among the live cells. **(B-E)** The *ex vivo* differentiated BM-DCs were subjected to forskolin (Forsk) treated CM collected from white and darkly pigmented melanocytes for 24 hrs and analyzed as described in *(A).*

Melanocytes respond to several different stimuli and stressors with increased pigment production to protect cells from external and internal damage.^13^ To mimic *in vivo* stress conditions and consequently induce pigment production in the *in vitro* cell culture settings, melanocytes of different pigment levels were treated with Forskolin (Forsk), which stimulates melanocyte pigment production by elevating cAMP levels.^54^ Forskolin-treated media alone increased the cDC subpopulation and decreased the pDC subpopulation (Suppl. Fig.1A, bottom panel) but had no effect on the maturation of the pDC subset (Suppl. Fig. 1F) when compared to media control (Suppl. Fig. 1A, top panel; Suppl. Fig. 1C). Exposure of BMDCs to Forsk-treated CM of black and white melanocytes increased the cDC1 subpopulation, had no effect on the cDC2 subset and decreased the pDC subpopulation (Suppl. Fig. 1D, E) when compared to Forsk-treated media alone (Suppl. Fig.1A, bottom panel). Interestingly, Forskolin-induced pigment production in black melanocytes (melan-a and melan-Cc) slightly affected the cDC1 subpopulation, had no effect on the cDC2 subset after incubation of the BMDCs with Forsk-treated CM of black melanocytes when compared to Forsk-treated CM of white cells (melan-e1 and melan-C) (Suppl. Fig. 1D, E). However, the number of pDCs was greatly reduced after BMDCs were exposed to the CM of Forsk-treated black melan-a melanocyte when compared to that of white melan-e1 cells (Suppl. Fig. 1D), but not after incubation with the CM of black melan-Cc cells (Suppl. Fig. 1E). Most importantly, pDCs exhibited an activated phenotype characterized by the significant up-regulation of CD40 (Fig. 1B), CD80 (Fig. 1C), CD86 (Fig. 1D) and MHC class II (Fig. 1E) cell surface markers when incubated with Forsk-treated CM of black melanocytes compared to Forsk-treated CM of white melanocytes. Like in the baseline conditions, forskolin treatment did not lead to significant difference in the level of cDC maturation following incubation with CM of black and white melanocytes (data not shown).

Together, these results suggest that dark pigmentation skews pDCs towards maturation.

### Dark pigmented melanocytes produce DC activating pro-inflammatory cytokines

Melanocytes are best known for producing pigment – melanin, thus taking central role in photoprotection against solar radiation.^55^ But melanocytes are also capable of secreting pro-inflammatory cytokines, consequently contributing to local immune responses.^26,27^ To address the question how black melanocytes trigger the maturation of DCs, the transcriptional expression of pro-inflammatory cytokines IL-6 and TNF-a, which are also recognized as inducers of DC maturation,^53,56^ was examined by qRT-PCR in un-stimulated black and white melanocytes. Figure 2A, B show that at baseline level, in an un-stimulated state black melanocytes express substantially more IL-6 and TNF-a mRNA when compared to white cells.

**Figure 2.**
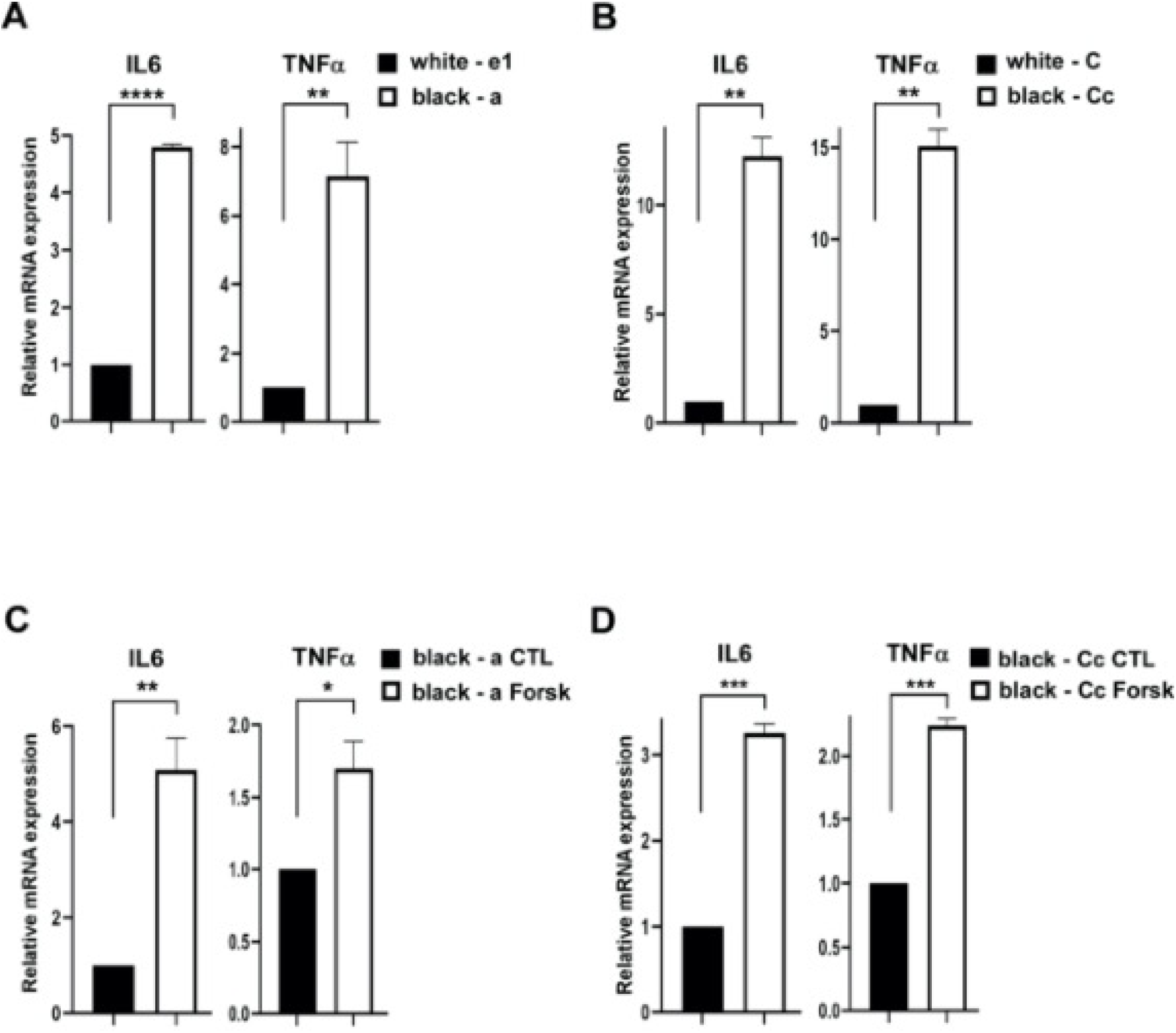
Dark pigment production is associated with elevated levels of DC-stimulating pro-inflammatory cytokines. **(A-B)** The transcriptional expression of IL-6, TNF-a was measured by qRT-PCR in white and black melanocytes. Graphs show quantification as fold of mRNA expression levels compared to control (white cells), mean□±□SD. **(C-D)** IL-6, TNF-a mRNA expression was assessed by qRT-PCR in control DMSO or Forsk treated darkly pigmented melanocytes. Graphs show quantification as fold of mRNA expression levels compared to control (DMSO), mean□±□SD. Statistical analysis was performed by unpaired, two-tailed *t* test with Welch’s correction: \**P*<0.05, \*\**P*<0.01, \*\*\**P*<0.001, \*\*\*\**P*<0.0001.

In addition, production of the inflammatory mediators was measured following activation of melanogenesis by Forsk treatment for 24 hours in darkly pigmented melanocytes. A significant increase in the mRNA expression of IL-6 (3-5 fold) and TNF-a (1.5-2.5 fold) was observed in Forsk stimulated black melanocytes compared to control DMSO treated black cells (Fig. 2C, D).

These results imply that the differences in the cytokine profiles of black and white melanocytes are potentially a contributing factor to the capacity of dark melanocytes to induce DC maturation.

### Melanocyte-secreted ECM protein, FMOD is associated with the immature phenotype of BMDCs

To further dissect the underlying mechanism for the differences observed between black and white melanocytes to induce the maturation of DCs, we turned our attention to a melanocyte-related ECM protein, FMOD. Our laboratory has previously demonstrated that depigmented (white) melanocytes secrete high levels of FMOD.^44^ To investigate the role of FMOD in pigment-associated DC maturation, we knocked down FMOD in white melanocytes, thereby mimicking the low levels of FMOD observed in darkly pigmented melanocytes.^44^

Stimulation of the BMDCs was accomplished by incubation with CMs collected from control and FMOD-KD melanocytes for 24 hrs. Flow cytometric analysis showed that a decrease in FMOD levels did not affect the distribution of the DC subpopulations (Suppl. Fig. 2) but correlated with a more mature phenotype of pDCs as determined by the higher surface expression of CD40, CD80, CD86 (Fig. 3A) and MHC class II (Fig. 3B) when compared to control.

**Figure 3.**
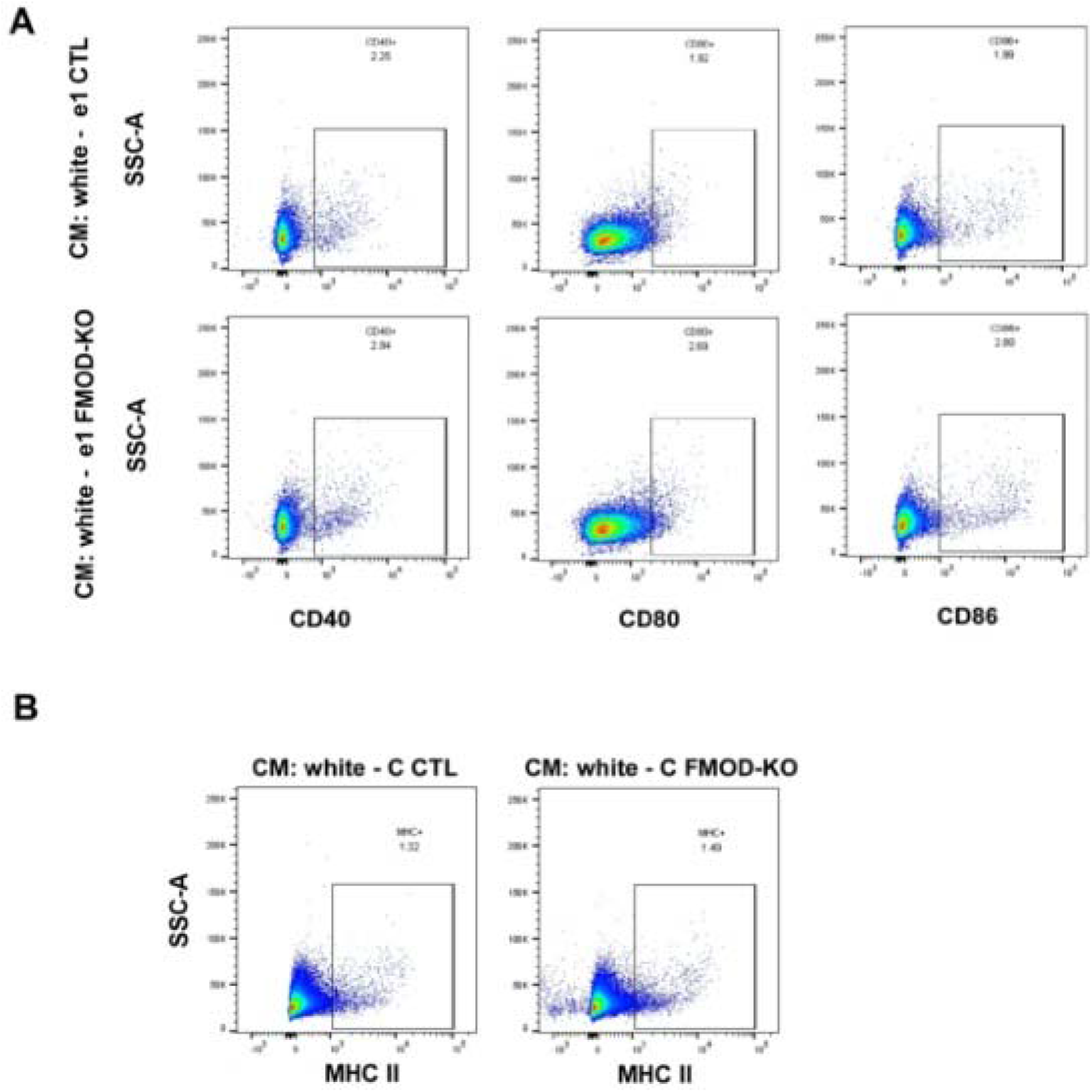
FMOD deletion promotes the maturation of pDCs. **(A-B)** Flt3L-BMDCs of 3 animals were incubated with CM derived from white control and FMOD-KD melanocytes. Twenty-four hours later the BMDCs were analyzed for the maturity of the pDCs by flow cytometry with CD40, CD80, CD86 **(A)** and MHC II **(B)** surface markers. Representative data are shown as dot plots, and numbers represent the percentage of CD11b’XCR-1’PDCA-1^+^B220^+^ CD40^+^/CD80^+^/CD86^+^/MHC II^+^ pDCs among the live cells.

In addition, the mRNA level of IL-6 and TNF-a was assessed and found to be elevated in the absence of FMOD (Fig. 4A, B), suggesting that FMOD influences the cytokine expression of melanocytes. Our previous work established that FMOD up-regulates TGF-β1 secretion.^44^ TGF-β1 appears to have a central role in maintaining DCs in an immature state.^47,57^ We observed a decrease in TGF-β1 mRNA expression in FMOD-KD white melanocytes (Fig. 4A, B) underscoring a potential role for FMOD in impacting DC maturation via TGF-β1.

**Figure 4.**
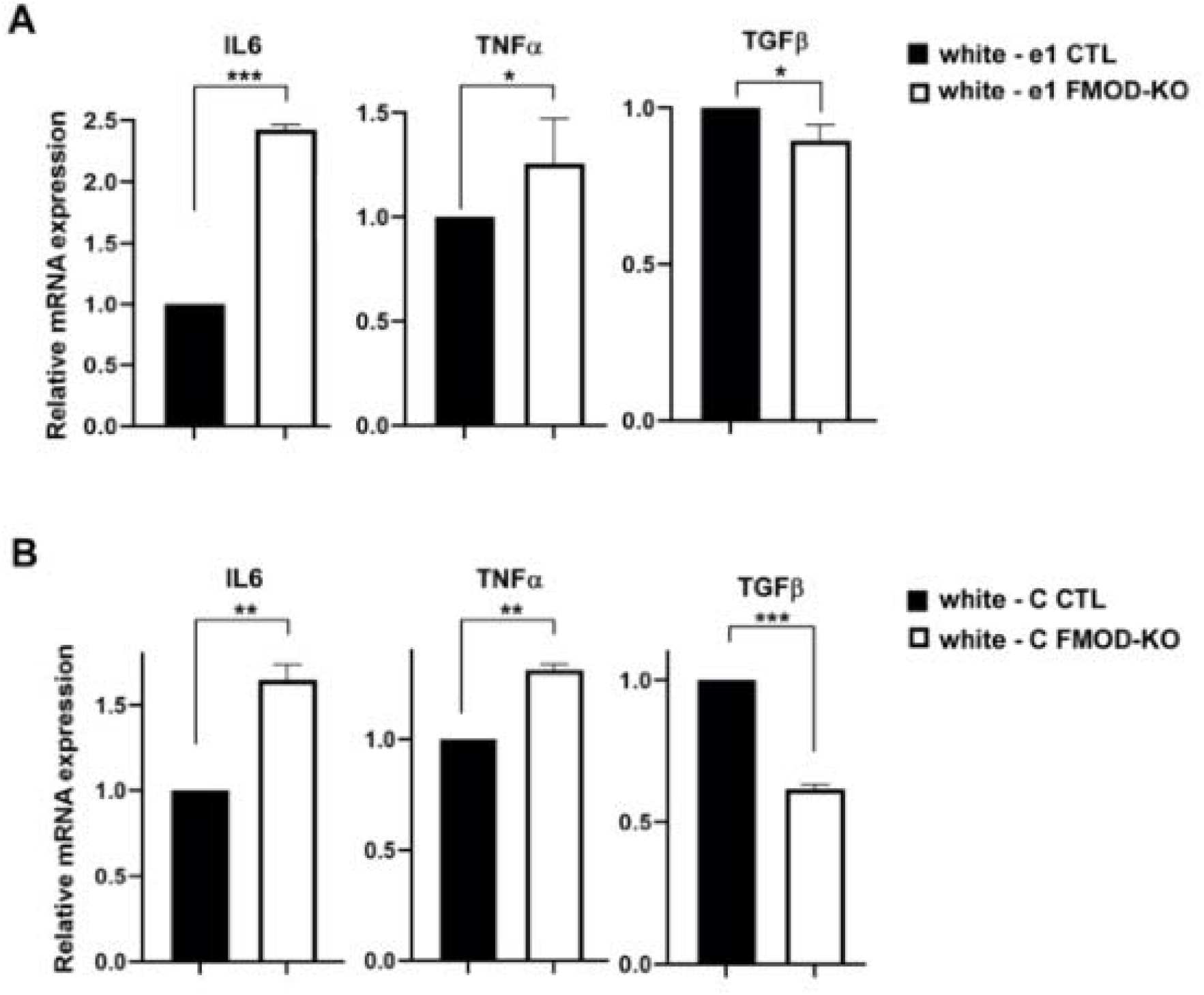
FMOD moderates the production of DC activating pro-inflammatory cytokines. **(A-B)** The transcriptional expression of IL-6, TNF-a and TGF-b1 was measured by qRT-PCR in white control and FMOD-KD melanocytes. Graphs show quantification as fold of mRNA expression levels compared to control, mean□±□SD. Statistical analysis was calculated by unpaired, two-tailed *t* test with Welch’s correction: \**P*<0.05, \*\**P*<0.01, \*\*\**P*<0.001.

Together these results point to the fact that white melanocytes because high level of FMOD interfere with DC maturation.

### Dark pigmentation and FMOD deletion correlate with elevated IL-3 levels

Recent work described IL-3 as an important factor not only for pDC survival^58^ but for its activation as well.^59^ Because we noted great differences in pDC maturation as marked by the elevated expression of cell surface maturation markers after incubation with CM of differentially pigmented melanocytes and with CM collected from melanocytes with FMOD-KD, we asked whether melanocytes produce different levels of IL-3.

First, we assessed the transcriptional expression of IL3 in un-stimulated black and white melanocytes using qRT-PCR. As shown on Fig 5A, elevated expression of IL3 was detected at base line in darkly pigmented melanocytes when compared to white melanocytes. Dependent on culture conditions, dark pigmented melanocytes can make various level of pigment, which can be confirmed by visual observation of the cells (Fig. 5B). Therefore, we examined the mRNA expression of IL3 in gray and black melan-Cc cells and found that dark pigmentation correlates with higher IL3 expression in comparison with light and no pigmentation (Fig. 5B).

**Figure 5.**
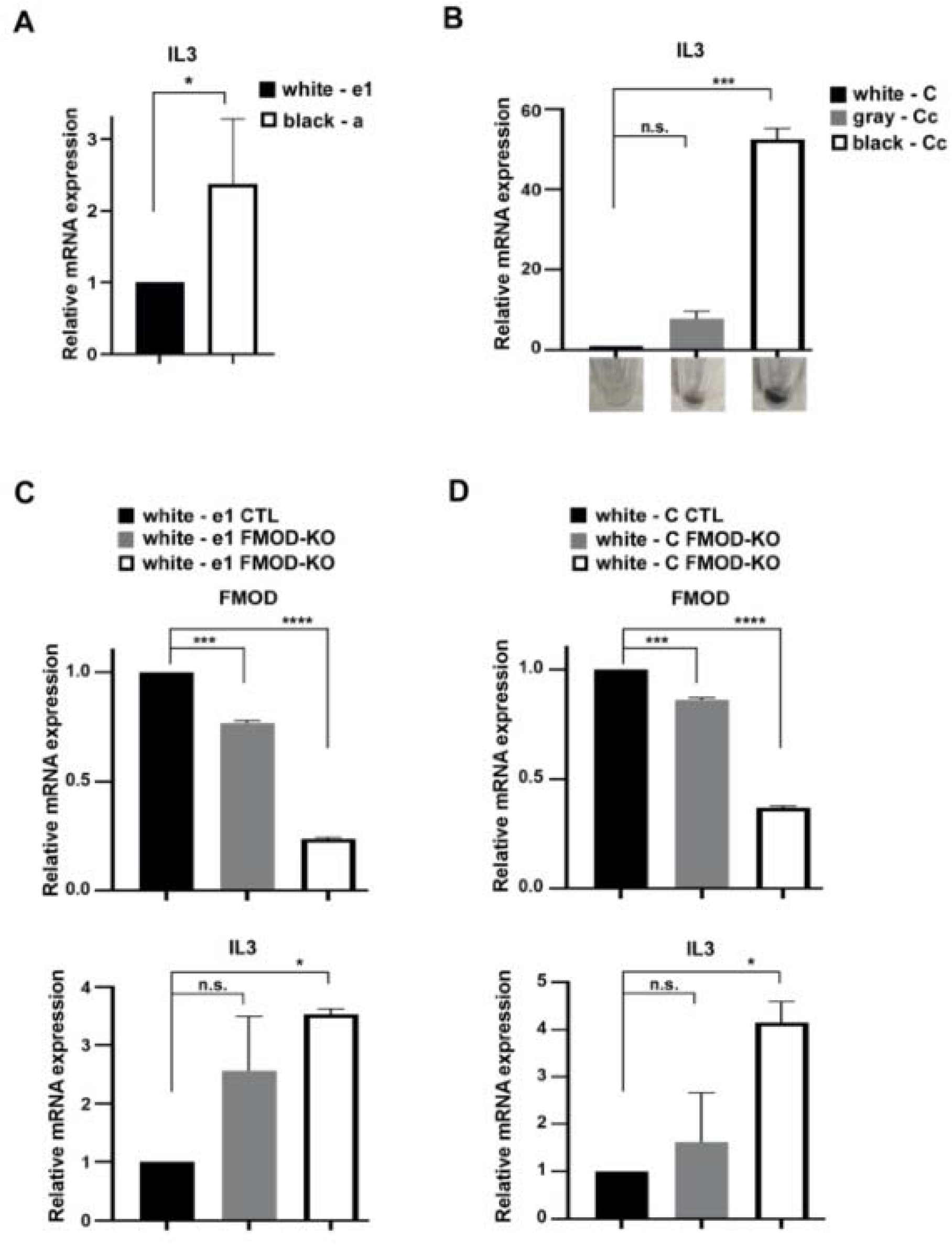
Dark pigmentation and FMOD depletion result in high IL-3 levels. **(A-B)** The mRNA expression of IL-3 was assessed by qRT-PCR in white and black or gray melanocytes. Graphs show quantification as fold of mRNA expression levels compared to control (white cells), mean□±□ SD. **(C-D)** IL-3 mRNA expression was measured by qRT-PCR in white control and FMOD-KD melanocytes. Graphs show quantification as fold of mRNA expression levels compared to control, mean□±□ SD. Statistical analysis was performed by unpaired, twotailed *t* test with Welch’s correction or by Ordinary One-Way ANOVA followed by Tukey’s multiple comparison posttest.: \**P*<0.05, \*\*\**P*<0.001, \*\*\*\**P*<0.0001.

Low pigment correlates with high FMOD^44^ and less pDC maturation (Fig. 3). We also measured IL3 mRNA expression in FMOD-KD white melanocytes to recapitulate the low expression of FMOD in black melanocytes and to further examine the potential role of FMOD in dark pigment-related pDC maturation. FMOD expression was confirmed by qRT-PCR as shown on Fig 5C, D. Interestingly, high mRNA levels of IL3 were detected in the absence of FMOD implicating FMOD in the transcriptional regulation of IL3 (Fig. 5C, D).

These results underscore the fact that dark pigmentation associated with low FMOD levels potentially triggers DC maturation as a result of increased IL3 expression.

## DISCUSSION

Epidemiological studies indicate that non-Hispanic African Americans are more likely to suffer from autoimmune disease, suggesting strong correlation between skin pigmentation and susceptibility to a particular autoimmune disease. With this study we investigated the possible underlying mechanisms for the ethnicity-dependent differences. We hypothesized that melanocyte pigmentation and melanocyte secreted FMOD by modulating DC maturation are contributing factors for the observed racial disparities.

To study the effect of melanocyte pigmentation on DC activity, we utilized Flt3L differentiated immature BMDCs incubated with CM of white and black murine epidermal melanocytes. Under these unstimulated, baseline conditions black melanocytes induced the maturation of the pDC subset of the BMDCs as shown by the elevated expressions of CD40, CD80, CD86 and MHC II on the DC surfaces. This finding was the very first indicator of potential differences in the baseline melanogenic activities of unstimulated differentially pigmented murine epidermal melanocytes. Of note, we did not observe differences in the degree of maturation of the cDC subset under the described culture conditions, so our focus was on the pDC subpopulation for the study. As expected, stimulation of inherent melanogenesis with forskolin in melanocytes led to an appreciable increase in the expression of the maturation markers on the pDCs incubated with Forsk-treated CM of black melanocytes when compared to that of white melanocytes. From these findings we can contemplate that the baseline darker pigmentation and subsequent induced pigment production of black epidermal melanocytes facilitate DC maturation underscoring the premise that higher melanin content is partially the source of differential immunity in individuals with dark vs. light skin complexion in response to various internal and external stimuli.

DC maturation can be triggered by numerous factors amongst them are pro-inflammatory cytokines.^53,56^ Cultured normal human epidermal melanocytes have been reported to release IL-6 and TNF-a after stimulation with bacterial LPS.^26^ The melanocyte secreted cytokines can act autocrine, paracrine and endocrine manner^60^, consequently melanocytes can contribute to an immune response locally.^26^ Abnormal cytokine production by melanocytes has been proposed as a contributing factor to changes in pigmentation and human diseases.^27^ We examined the transcriptional expression of IL-6 and TNF-□ and found that black melanocytes expressed significantly higher mRNA levels of the pro-inflammatory cytokines not only at baseline but also after stimulation of melanogenesis with Forsk. Most importantly, our result implies that the intrinsic dark pigmented melanocytes via the increased expression of pro-inflammatory mediators compel DCs to mature. To the best of our knowledge, this is the first study to show that even the baseline melanogenic activity of differently pigmented epidermal melanocytes affects pDC maturation. It is tempting to speculate that darker skin is a high-risk factor for a pro-inflammatory microenvironment and therefore for the development of inflammation related diseases as a result of more pronounced DC maturation.

Our early work showed that low pigment production correlates with high levels of FMOD, a melanocyte-secreted ECM protein.^61^ Because dark pigmentation stimulated pDC maturation, we hypothesized that by decreasing the level of pigment-related FMOD in white cells we can recapitulate the maturation effect of dark pigment on pDCs in white cells. As expected, we observed that FMOD-KD in white cells resulted in increased pDC maturation and proinflammatory cytokine expression. This suggests a potential role for FMOD in pigment-induced DC maturation.

pDCs are a rare type of dendritic cell subset that typically originate from the bone marrow and exhibit plasma cell morphology.^62^ First and foremost, pDCs are specialized in the secretion of high amounts of type I IFNs upon encounter with viruses, but they have been associated with autoimmune diseases as well. Peripheral pDCs and cDCs have been observed to decrease in patients with autoimmune diseases; however, in the autoimmune lesions the number of pDCs have increased. Over the course of their life cycle, pDCs stop producing cytokines and acquire the ability for antigen (Ag) presentation via MHC molecules and for subsequent T cell activation.^35,62^ Ag presentation is accompanied by the upregulation of costimulatory receptor molecules CD40, CD80, CD83, and CD86 on DC cell membranes.^53^ pDCs primarily are activated when their endosomal Toll-like receptor (TLR)7 or TLR9 get ligated.^36^ IL-3 is known to support pDC growth and survival, but a recent report also identified it as a pDC activator.^58,59^ Since dark pigment melanocytes stimulate pDC maturation, we examined the production of IL-3 in the differentially pigmented melanocytes. We observed that the transcriptional expression of IL-3 is correlative with the intensity of pigmentation and level of FMOD. These data provide a novel mechanistic basis for stimulating pDC maturation through IL-3 produced by dark pigmented melanocytes with low FMOD levels.

Though, as weak APCs melanocytes themselves can present to T cells (XX) but based on our results it is tempting to consider dark pigmented melanocytes as signal amplifiers for DC activation and maturation by promoting an inflammatory environment via cytokine secretion. Hyperpigmentation of the skin following inflammation is a widely recognized fact in clinical settings^26^ suggesting that melanocyte biology and skin pigmentation is altered during inflammatory states. Our work implies that darker skin is a higher risk factor for a stronger inflammatory response. Consequently, we postulate the existence of a self-escalating positive feedback loop, wherein inflammation correlates with darker pigmentation and higher melanin content results in more inflammation.

Here, we utilized epidermal melanocytes as our model system, and it is tempting to speculate that the findings of this study hold far reaching consequences. Melanocytes are situated in the epidermis, hair bulbs, eyes, ears, heart, and brain,^8,9^ and able to secret pro-inflammatory cytokines, enabling them to affect tissue microenvironment throughout the body, and to modulate not only local but systemic immune responses.

## ACKNOWLEDGMENT

This study was supported in part by a grant from NIH (RO1EY024046).

## SUPPLEMENTAL

**Supplementary Figure 1.**
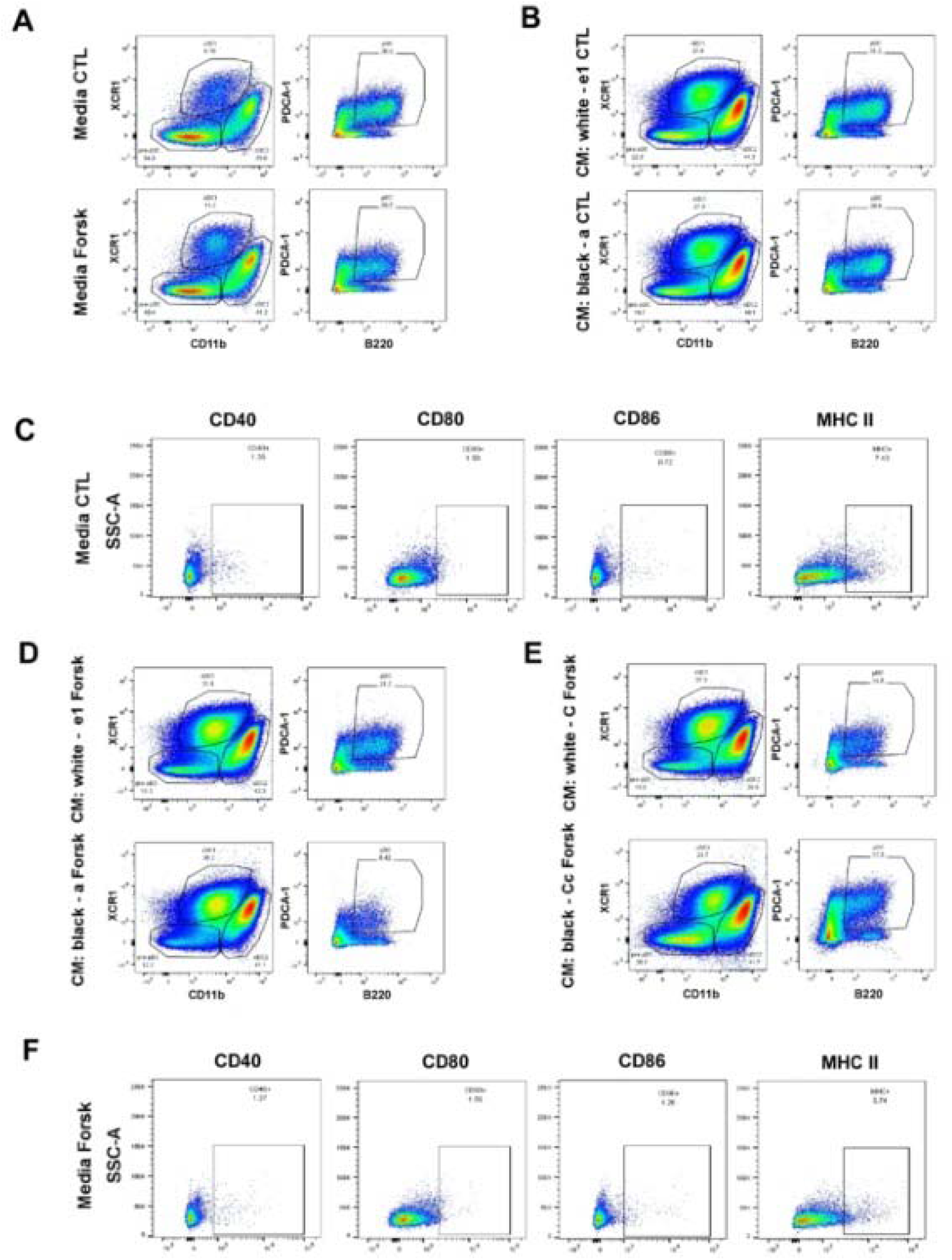
**(A)** Mouse Flt3L differentiated BMDCs of 3 animals were incubated with control DMSO and Forsk treated media. Twenty-four hours later the BMDCs were analyzed by flow cytometry. Dot plots show the gating strategy of the DC subpopulations including cDC1 (CD11b^-^XCR-1^+^), cDC2 (CD11b^+^XCR-1^-^) and pDC (CD11b^-^XCR-1^-^PDCA-1^+^B220^+^) among the live cells. **(B)** Flt3L-BMDCs were exposed to CM derived from control DMSO treated white (melan e1) and black (melan a) melanocytes and analyzed as described in *(A).* **(C)** BMDCs described in (*A*) were analyzed for the maturity of the pDC subpopulation by flow cytometry with CD40, CD80, CD86 and MHC II surface markers. Representative data are shown as dot plots, and numbers represent the percentage of CD11b^-^XCR-1^-^PDCA-1^+^B220^+^ CD40^+^/CD80^+^/CD86^+^/MHC II^+^ pDCs among the live cells. **(D-E)** Flt3L-BMDCs were subjected to Forsk treated CM collected from white and black melanocytes and analyzed as described in (*A)*. **(F)** BMDCs described in (*A*) were analyzed for the maturity of the pDC subpopulation by flow cytometry as described in (*C*).

**Supplementary Figure 2.**
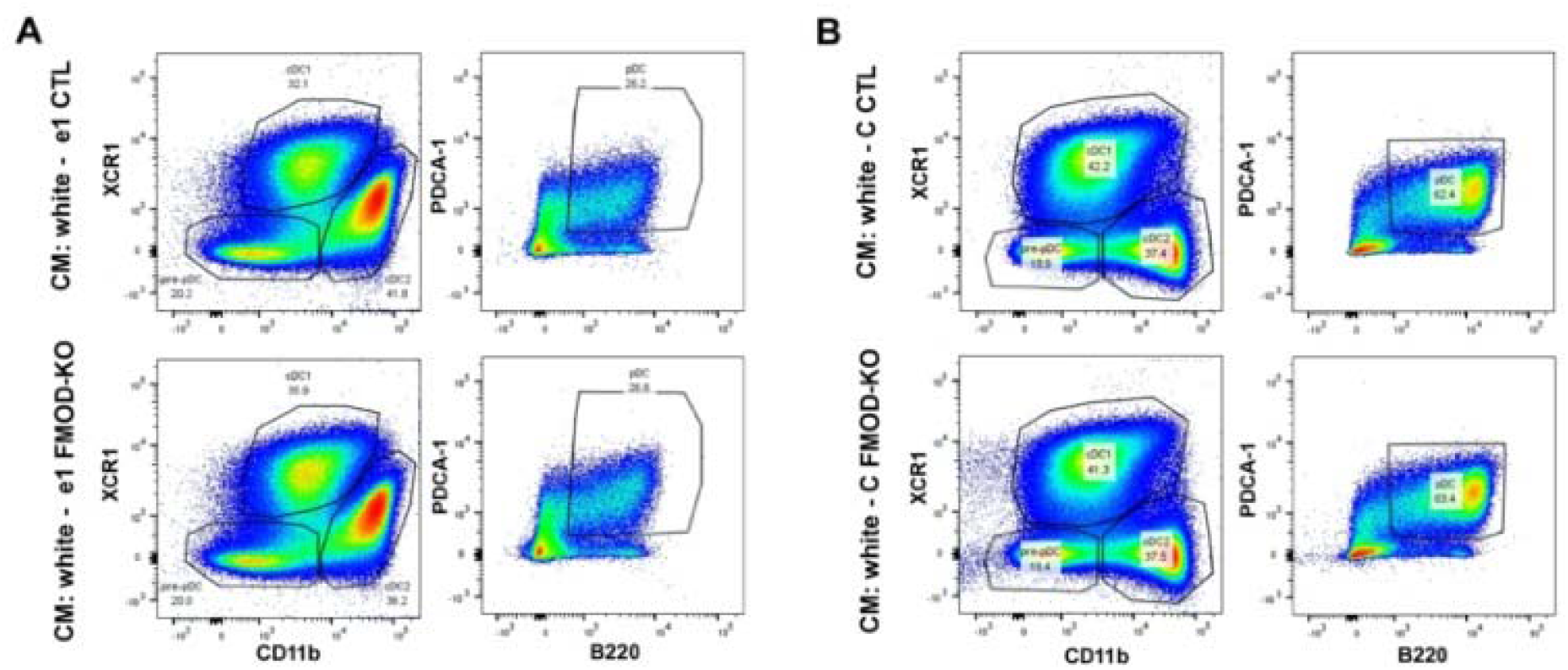
**(A-B)** Flt3L-BMDCs of 3 animals were incubated with CM derived from white control and FMOD-KD melanocytes. Twenty-four hours later the BM-DCs were analyzed by flow cytometry. Dot plots show the gating strategy of the DC subpopulations including cDC1 (CD11b^-^XCR-1^+^), cDC2 (CD11b^+^XCR-1^-^) and pDC (CD11b^-^XCR-1^-^PDCA-1^+^B220^+^) among the live cells.

## REFERENCES

1. Mealy, M.A., et al. Mortality in neuromyelitis optica is strongly associated with African ancestry. Neurol Neuroimmunol Neuroinflamm 5, e468 (2018).

2. Chauhan, K., Scaife, S. & Rosenbaum, J.T. Uveitis and health disparities: results from the National Inpatient Sample. Br J Ophthalmol 103, 1301–1305 (2019).

3. Shaw, T.E., Currie, G.P., Koudelka, C.W. & Simpson, E.L. Eczema prevalence in the United States: data from the 2003 National Survey of Children’s Health. J Invest Dermatol 131, 67–73 (2011).

4. Pons-Estel, G.J., Alarcon, G.S., Scofield, L., Reinlib, L. & Cooper, G.S. Understanding the epidemiology and progression of systemic lupus erythematosus. Semin Arthritis Rheum 39, 257–268 (2010).

5. Somers, E.C., et al. Population-based incidence and prevalence of systemic lupus erythematosus: the Michigan Lupus Epidemiology and Surveillance program. Arthritis Rheumatol 66, 369–378 (2014).

6. Szabo, G. The number of melanocytes in human epidermis. Br Med J 1, 1016–1017 (1954).

7. Mort, R.L., Jackson, I.J. & Patton, E.E. The melanocyte lineage in development and disease. Development 142, 620–632 (2015).

8. Yajima, I. & Larue, L. The location of heart melanocytes is specified and the level of pigmentation in the heart may correlate with coat color. Pigment Cell Melanoma Res 21, 471–476 (2008).

9. Gudjohnsen, S.A., et al. Meningeal Melanocytes in the Mouse: Distribution and Dependence on Mitf. Front Neuroanat 9, 149 (2015).

10. Yaron, I., Zakheim, A.R., Oluwole, S.F. & Hardy, M.A. Effects of ultraviolet B irradiation on human natural killer cell and lymphokine activated killer cell activity: therapeutic potential in bone marrow transplantation and tumor immunotherapy. Transplant Proc 27, 1379 (1995).

11. Swope, V.B., Abdel-Malek, Z., Kassem, L.M. & Nordlund, J.J. Interleukins 1 alpha and 6 and tumor necrosis factor-alpha are paracrine inhibitors of human melanocyte proliferation and melanogenesis. J Invest Dermatol 96, 180–185 (1991).

12. Slominski, A., et al. A novel pathway for sequential transformation of 7-dehydrocholesterol and expression of the P450scc system in mammalian skin. Eur J Biochem 271, 4178–4188 (2004).

13. D’Mello, S.A., Finlay, G.J., Baguley, B.C. & Askarian-Amiri, M.E. Signaling Pathways in Melanogenesis. Int J Mol Sci 17(2016).

14. Preising, M.N., Forster, H., Gonser, M. & Lorenz, B. Screening of TYR, OCA2, GPR143, and MC1R in patients with congenital nystagmus, macular hypoplasia, and fundus hypopigmentation indicating albinism. Mol Vis 17, 939–948 (2011).

15. van Tuyn, J., et al. Oncogene-Expressing Senescent Melanocytes Up-Regulate MHC Class II, a Candidate Melanoma Suppressor Function. J Invest Dermatol 137, 2197–2207 (2017).

16. Cichorek, M., Wachulska, M., Stasiewicz, A. & Tyminska, A. Skin melanocytes: biology and development. Postepy Dermatol Alergol 30, 30–41 (2013).

17. Lu, Y., Zhu, W.Y., Tan, C., Yu, G.H. & Gu, J.X. Melanocytes are potential immunocompetent cells: evidence from recognition of immunological characteristics of cultured human melanocytes. Pigment Cell Res 15, 454–460 (2002).

18. Le Poole, I.C., et al. Phagocytosis by normal human melanocytes in vitro. Exp Cell Res 205, 388–395 (1993).

19. Le Poole, I.C., et al. A novel, antigen-presenting function of melanocytes and its possible relationship to hypopigmentary disorders. J Immunol 151, 7284–7292 (1993).

20. Hari, A., Flach, T.L., Shi, Y. & Mydlarski, P.R. Toll-like receptors: role in dermatological disease. Mediators Inflamm 2010, 437246 (2010).

21. Gasque, P. & Jaffar-Bandjee, M.C. The immunology and inflammatory responses of human melanocytes in infectious diseases. J Infect 71, 413–421 (2015).

22. Huang, D.J., Chen, R.F. & Wang, B.Y. [Catheter ablation in eight patients with Wolff-Parkinson-White syndrome]. Zhonghua Xin Xue Guan Bing Za Zhi 17, 90–92, 126 (1989).

23. Zachariae, C.O., Thestrup-Pedersen, K. & Matsushima, K. Expression and secretion of leukocyte chemotactic cytokines by normal human melanocytes and melanoma cells. J Invest Dermatol 97, 593–599 (1991).

24. Slominski, A., Wortsman, J., Luger, T., Paus, R. & Solomon, S. Corticotropin releasing hormone and proopiomelanocortin involvement in the cutaneous response to stress. Physiol Rev 80, 979–1020 (2000).

25. Swope, V.B., et al. Synthesis of interleukin-1 alpha and beta by normal human melanocytes. J Invest Dermatol 102, 749–753 (1994).

26. Tam, I., Dzierzega-Lecznar, A. & Stepien, K. Differential expression of inflammatory cytokines and chemokines in lipopolysaccharide-stimulated melanocytes from lightly and darkly pigmented skin. Exp Dermatol 28, 551–560 (2019).

27. Tam, I. & Stepien, K. Secretion of proinflammatory cytokines by normal human melanocytes in response to lipopolysaccharide. Acta Biochim Pol 58, 507–511 (2011).

28. Banchereau, J. & Steinman, R.M. Dendritic cells and the control of immunity. Nature 392, 245–252 (1998).

29. Song, M.G., et al. Role of aquaporin 3 in development, subtypes and activation of dendritic cells. Mol Immunol 49, 28–37 (2011).

30. Geissmann, F., et al. Development of monocytes, macrophages, and dendritic cells. Science 327, 656–661 (2010).

31. Merad, M., Sathe, P., Helft, J., Miller, J. & Mortha, A. The dendritic cell lineage: ontogeny and function of dendritic cells and their subsets in the steady state and the inflamed setting. Annu Rev Immunol 31, 563–604 (2013).

32. Sichien, D., Lambrecht, B.N., Guilliams, M. & Scott, C.L. Development of conventional dendritic cells: from common bone marrow progenitors to multiple subsets in peripheral tissues. Mucosal Immunol 10, 831–844 (2017).

33. Shortman, K., Sathe, P., Vremec, D., Naik, S. & O’Keeffe, M. Plasmacytoid dendritic cell development. Adv Immunol 120, 105–126 (2013).

34. Bekeredjian-Ding, I., Greil, J., Ammann, S. & Parcina, M. Plasmacytoid Dendritic Cells: Neglected Regulators of the Immune Response to Staphylococcus aureus. Front Immunol 5, 238 (2014).

35. Swiecki, M. & Colonna, M. The multifaceted biology of plasmacytoid dendritic cells. Nat Rev Immunol 15, 471–485 (2015).

36. Gilliet, M., Cao, W. & Liu, Y.J. Plasmacytoid dendritic cells: sensing nucleic acids in viral infection and autoimmune diseases. Nat Rev Immunol 8, 594–606 (2008).

37. O’Keeffe, M., et al. Dendritic cell precursor populations of mouse blood: identification of the murine homologues of human blood plasmacytoid pre-DC2 and CD11c+ DC1 precursors. Blood 101, 1453–1459 (2003).

38. Iozzo, R.V. & Schaefer, L. Proteoglycan form and function: A comprehensive nomenclature of proteoglycans. Matrix Biol 42, 11–55 (2015).

39. Mikaelsson, E., et al. Fibromodulin, an extracellular matrix protein: characterization of its unique gene and protein expression in B-cell chronic lymphocytic leukemia and mantle cell lymphoma. Blood 105, 4828–4835 (2005).

40. Lee, Y.H. & Schiemann, W.P. Fibromodulin suppresses nuclear factor-kappaB activity by inducing the delayed degradation of IKBA via a JNK-dependent pathway coupled to fibroblast apoptosis. J Biol Chem 286, 6414–6422 (2011).

41. Lee, E.J., et al. Fibromodulin: a master regulator of myostatin controlling progression of satellite cells through a myogenic program. FASEB J 30, 2708–2719 (2016).

42. Zheng, Z., et al. Reprogramming of human fibroblasts into multipotent cells with a single ECM proteoglycan, fibromodulin. Biomaterials 33, 5821–5831 (2012).

43. Zheng, Z., et al. Fibromodulin Enhances Angiogenesis during Cutaneous Wound Healing. Plast Reconstr Surg Glob Open 2, e275 (2014).

44. Adini, I., et al. Melanocyte-secreted fibromodulin promotes an angiogenic microenvironment. J Clin Invest 124, 425–436 (2014).

45. Jan, A.T., Lee, E.J. & Choi, I. Fibromodulin: A regulatory molecule maintaining cellular architecture for normal cellular function. Int J Biochem Cell Biol 80, 66–70 (2016).

46. Zeng-Brouwers, J., Pandey, S., Trebicka, J., Wygrecka, M. & Schaefer, L. Communications via the Small Leucine-rich Proteoglycans: Molecular Specificity in Inflammation and Autoimmune Diseases. J Histochem Cytochem 68, 887–906 (2020).

47. Kel, J.M., Girard-Madoux, M.J., Reizis, B. & Clausen, B.E. TGF-beta is required to maintain the pool of immature Langerhans cells in the epidermis. J Immunol 185, 3248–3255 (2010).

48. Alexeev, V. & Yoon, K. Gene correction by RNA-DNA oligonucleotides. Pigment Cell Res 13, 72–79 (2000).

49. Bennett, D.C., Cooper, P.J. & Hart, I.R. A line of non-tumorigenic mouse melanocytes, syngeneic with the B16 melanoma and requiring a tumour promoter for growth. Int J Cancer 39, 414–418 (1987).

50. Hida, T., et al. Agouti protein, mahogunin, and attractin in pheomelanogenesis and melanoblast-like alteration of melanocytes: a cAMP-independent pathway. Pigment Cell Melanoma Res 22, 623–634 (2009).

51. Naik, S.H., O’Keeffe, M., Proietto, A., Shortman, H.H. & Wu, L. CD8+, CD8-, and plasmacytoid dendritic cell generation in vitro using flt3 ligand. Methods Mol Biol 595, 167–176 (2010).

52. Iveson-Iveson, J. Haloperidol; a useful psychiatric drug. Nurs Mirror Midwives J 143, 77 (1976).

53. Mbongue, J.C., Nieves, H.A., Torrez, T.W. & Langridge, W.H. The Role of Dendritic Cell Maturation in the Induction of Insulin-Dependent Diabetes Mellitus. Front Immunol 8, 327 (2017).

54. Amaro-Ortiz, A., Vanover, J.C., Scott, T.L. & D’Orazio, J.A. Pharmacologic induction of epidermal melanin and protection against sunburn in a humanized mouse model. J Vis Exp (2013).

55. Slominski, A., Tobin, D.J., Shibahara, S. & Wortsman, J. Melanin pigmentation in mammalian skin and its hormonal regulation. Physiol Rev 84, 1155–1228 (2004).

56. Brasel, K., De Smedt, T., Smith, J.L. & Maliszewski, C.R. Generation of murine dendritic cells from flt3-ligand-supplemented bone marrow cultures. Blood 96, 3029–3039 (2000).

57. Yamaguchi, Y., Tsumura, H., Miwa, M. & Inaba, K. Contrasting effects of TGF-beta 1 and TNF-alpha on the development of dendritic cells from progenitors in mouse bone marrow. Stem Cells 15, 144–153 (1997).

58. Lutz, M.B. IL-3 in dendritic cell development and function: a comparison with GM-CSF and IL-4. Immunobiology 209, 79–87 (2004).

59. Grzes, K.M., et al. Plasmacytoid dendritic cell activation is dependent on coordinated expression of distinct amino acid transporters. Immunity 54, 2514–2530 e2517 (2021).

60. Imokawa, G. Autocrine and paracrine regulation of melanocytes in human skin and in pigmentary disorders. Pigment Cell Res 17, 96–110 (2004).

61. Adini, I., et al. Melanocyte pigmentation inversely correlates with MCP-1 production and angiogenesis-inducing potential. FASEB J 29, 662–670 (2015).

62. Ye, Y., Gaugler, B., Mohty, M. & Malard, F. Plasmacytoid dendritic cell biology and its role in immune-mediated diseases. Clin Transl Immunology 9, e1139 (2020).

